# Quantifying the contribution of dominance effects to complex trait variation in biobank-scale data

**DOI:** 10.1101/2020.11.10.376897

**Authors:** Ali Pazokitoroudi, Alec M. Chiu, Kathryn S. Burch, Bogdan Pasaniuc, Sriram Sankararaman

**Affiliations:** Department of Computer Science, UCLA, Los Angeles, California; Bioinformatics Interdepartmental Program, UCLA, Los Angeles, California; Department of Pathology and Laboratory Medicine, David Geffen School of Medicine, UCLA, Los Angeles, California; Department of Human Genetics, David Geffen School of Medicine, UCLA, Los Angeles, California; Department of Computational Medicine, David Geffen School of Medicine, UCLA, Los Angeles, California

## Abstract

The proportion of variation in complex traits that can be attributed to non-additive genetic effects has been a topic of intense debate. The availability of Biobank-scale datasets of genotype and trait data from unrelated individuals opens up the possibility of obtaining precise estimates of the contribution of non-additive genetic effects. We present an efficient method that can partition the variation in complex traits into variance that can be attributed to additive (*additive heritability*) and dominance (*dominance heritability*) effects across all genotyped SNPs in a large collection of unrelated individuals. Over a wide range of genetic architectures, our method yields unbiased estimates of heritability. We applied our method, in turn, to array genotypes as well as imputed genotypes (at common SNPs with minor allele frequency, MAF > 1%) and 50 quantitative traits measured in 291, 273 unrelated white British individuals in the UK Biobank. Averaged across these 50 traits, we find that additive heritability on array SNPs is 21.86% while dominance heritability is 0.13% (about 0.48% of the additive heritability) with qualitatively similar results for imputed genotypes. We find no evidence for dominance heritability (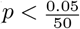 accounting for the number of traits tested) and estimate that dominance heritability is unlikely to exceed 1% for the traits analyzed. Our analyses indicate a limited contribution of dominance heritability to complex trait variation.

## Introduction

Variation in complex traits can be partitioned into variation due to additive, dominance and epistatic effects [1]. Despite decades of theoretical and experimental efforts, the quantification of non-additive genetic variation in outbred populations such as humans remains challenging [2, 3, 4, 5, 6]. One approach to estimate non-additive sources of heritability in humans have been focused on comparing phenotypic similarity between close relatives [7]. However, confounding with common environmental factors within families can lead to biased estimates. Further, the limited sample sizes of family and twin studies lead to large standard errors in estimates of non-additive effects. An alternative approach relies on the analysis of unrelated individuals. Using such an approach, Zhu *et al*. [8] estimated additive and dominance heritability at genotyped SNPs in 79 to find that dominance heritability was about a fifth of additive heritability on average although none of the traits were found to have significant dominance heritability.

The relatively small estimates of dominance heritability from prior studies [3, 8] suggest that achieving sufficient power to detect dominance heritability will require the analysis of large numbers of unrelated individuals and methods that can be run on these large sample sizes. We propose a variance components method that can jointly estimate the heritability due to additive and dominance effects attributed to SNPs genotyped across hundreds of thousands of individuals. Further, our method can jointly fit multiple additive and dominance variance components thereby allowing it to provide unbiased estimates of heritability for genetic architectures in which SNP effect sizes vary as a function of minor allele frequency (MAF) and linkage disequilibrium (LD).

Our method obtains unbiased estimates of additive and dominance heritability under a range of MAF and LD-dependent architectures while controlling the false positive rate of rejecting the null hypothesis of no dominance heritability under genetic architectures that assume no dominance. Applying our method to a total of 50 continuous traits measured in 291, 273 unrelated white British individuals in the UK Biobank, we find that additive heritability is 21.86% on average while dominance heritability is 0.13% on average (about 0.48% of the heritability attributed to additive effects) across common array SNPs (*M* = 459, 792 SNPs, MAF > 1%). Analyzing common imputed SNPs (*M* = 4, 824, 392, MAF > 1%), we find that additive heritability is 22.83% on average while dominance heritability is 0.06% on average (about 0.47% of the heritability attributed to additive effects). We find no evidence for traits that have non-zero dominance heritability after correcting for multiple testing 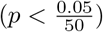. Based on the power estimates of our method, we estimate that dominance heritability is unlikely to exceed 1% for the traits analyzed.

## Results

### Methods overview

We aim to fit a variance components model that relates phenotypes ***y*** measured across *N* individuals to their additive and dominance values over *M* SNPs (while allowing for multiple additive and dominance components):

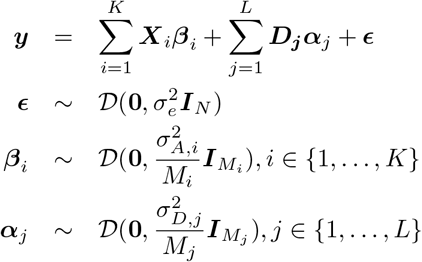

Here 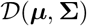 is an arbitrary distribution over a random vector with mean ***μ*** and covariance matrix **Σ**. Here SNPs are partitioned into *K* additive categories and *L* dominance categories. Here ***X**_i_* and ***D**_j_* are the *N* × *M_i_* and *N* × *M_j_* matrices consisting of standardized additive and dominance values of SNPs belonging to additive category *i* and dominance category *j* respectively, 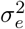 is the residual variance, and 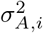 and 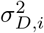 are the variance component of *i*-th additive and dominance categories respectively.

We encode additive and dominance effects using a representation that leads to uncorrelated variance components [8]. For alleles *A* and *B* at a SNP with the frequency of allele B denoted by *f_B_*, the additive *v_A_* and dominance *v_D_* values of the genotypes are defined as follows :

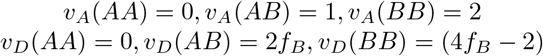

The variance explained by additive variation (*additive heritability*) at all SNPs is defined as:

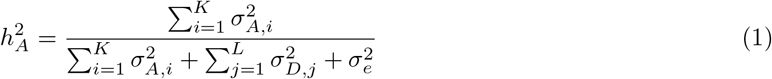

The variance explained by dominance variation (*dominance heritability*) at all SNPs is defined as:

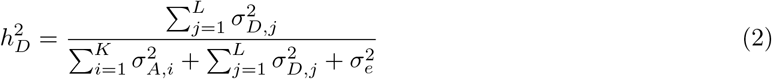

The proposed model extends previous models by introducing the component corresponding to dominance effects in addition to the additive effects [9]. Further, the proposed model allows for the joint estimation of additive and dominance components, e.g., corresponding to SNPs with varying minor allele frequency (MAF) and linkage disequilibrium (LD) annotations that have been previously shown to lead to relatively unbiased estimates of SNP heritability [10, 9].

The key inference problem in this model is the estimation of the variance components: 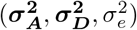 where 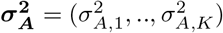 and 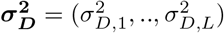. We use a scalable method-of-moments estimator, *i.e*., finding values of the variance components such that the population moments match the sample moments [11, 12, 13, 14, 15]. Our method uses a randomized algorithm that avoids explicitly computing genetic relatedness matrices. Instead, it operates on a smaller matrix formed by multiplying the input genotype matrix with a small number of random vectors allowing it to scale to large samples (Methods). We estimate standard errors (SE) using an efficient block Jackknife over SNPs with 100 blocks.

### Accuracy of estimates of dominance heritability in simulations

Previous studies estimate a relatively small contribution of dominance heritability for complex traits [8] so that we would like to test the false positive rate of a test of the hypothesis of no dominance heritability. To assess the false positive rate of our method, we performed simulations in the absence of dominance effects (*M* = 459, 792 SNPs, *N* = 291, 273 individuals). Since additive SNP effects tend to vary as a function of MAF and LD patterns at the SNP [10, 16] and SNP heritability estimates tend to be sensitive to these assumptions, we simulated phenotypes according to 16 MAF and LD-dependent architectures by varying the additive heritability, the proportion of variants that have non-zero effects (causal variants or CVs), the distribution of causal variants across minor allele frequencies (CVs distributed across all minor allele frequency bins or CVs restricted to either common or low-frequency bins), and the form of coupling between the SNP effect size and MAF as well as LD. The key parameter in applying our method is the number of random vectors *B* which we set to 10. To obtain unbiased estimates, we also do not constrain the estimates of the variance components (allowing for negative estimates).

Recent studies have shown that methods that fit a single additive variance component yield biased estimates of SNP heritability due to the LD and MAF dependent architecture of complex traits [10, 16, 17] while models that allow for SNP effects to vary with MAF and LD obtain relatively unbiased estimates [10, 16, 9]. Thus, we ran our method using 24 bins for additive effects (based on 6 MAF and 4 LD bins) and a single bin for dominance effects (although our method allows for fitting multiple dominance bins). Across the range of genetic architectures, we obtained accurate estimates of 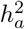 when we jointly fit additive and dominance heritability: biases range from −2 × 10^−3^ to 2 × 10^−3^ where 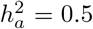 (Figure 1). We also obtain unbiased estimates of 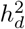 with biases ranging from −5 × 10^−5^ to 6 × 10^−4^ where 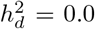 (Figure 1). Importantly, the false positive rate of rejecting the null hypothesis of no dominance heritability across 16 diverse genetic architecture is controlled at level 0. 05 (see Table 1). We performed additional simulations that demon-strateaccurate heritability estimates for a smaller sample size of *N* = 10, 000 individuals (Supplementary Figure S1 and Table S1).

**Figure 1:**
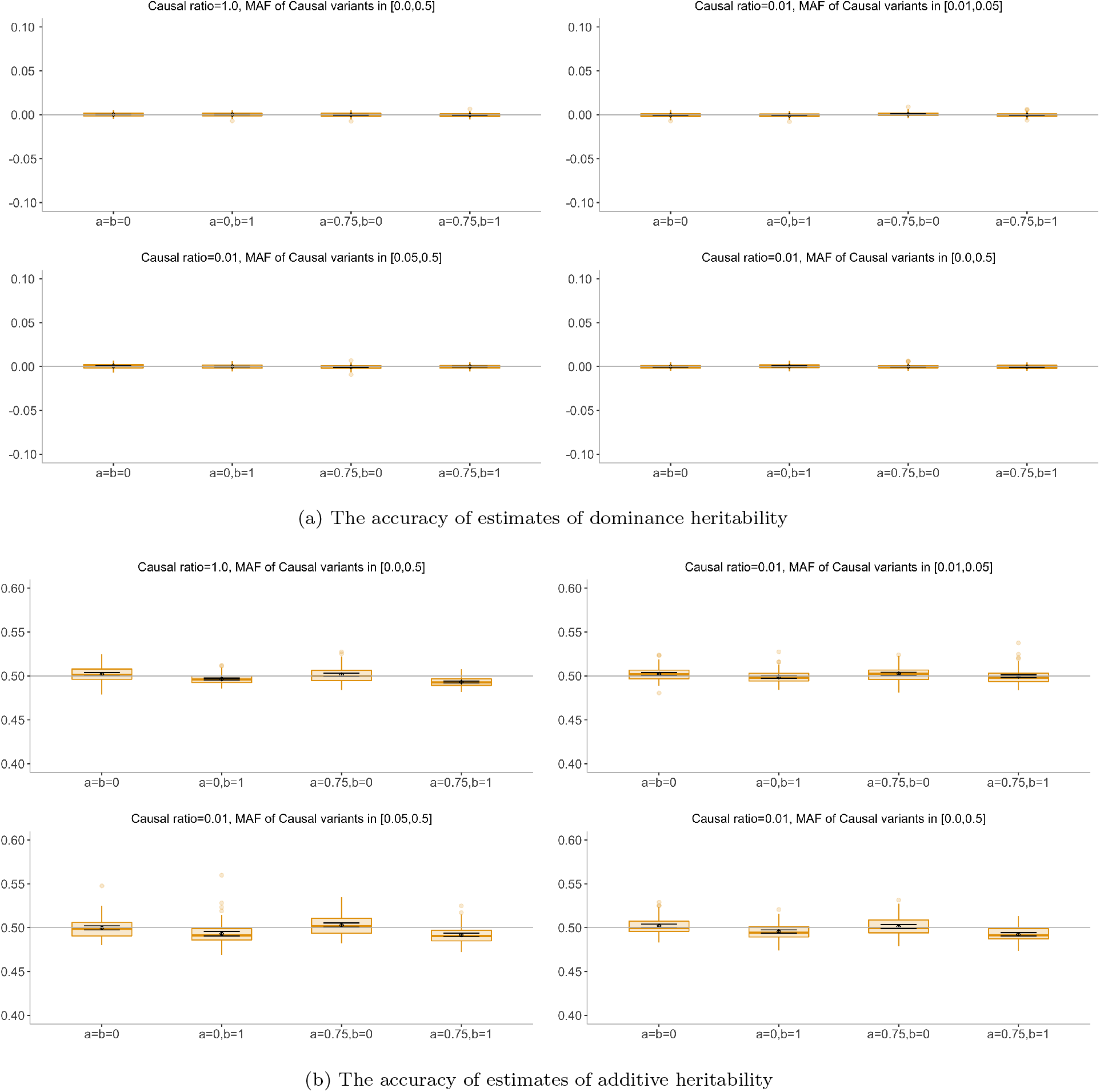
The accuracy of estimates of dominance and additive heritabilities in simulations with no dominance heritability (*N* = 291, 273 unrelated individuals, *M* = 459, 792 array SNPs). In **a** and **b**: We plot estimates from our method in the absence of dominance effect under 16 different genetic architectures. We varied the MAF range of causal variants (MAF of CV), the coupling of MAF with effect size (*a*), and the effect of local LD on effect size (*b* = 0 indicates no LDAK weights and *b* = 1 indicates LDAK weights (see Methods). We ran 100 replicates where the true additive and dominance heritabilities of the phenotype are 0.5 and 0.0 respectively. We ran our method using a single dominance bin and 24 additive bins formed by the combination of 6 bins based on MAF as well as 4 bins based on quartiles of the LDAK score of a SNP (see Methods). Black points and error bars represent the mean and ±2 SE. Each boxplot represents estimates from 100 simulations.

**Table 1:** Calibration of tests of dominance heritability. We assess the false positive rate of tests of dominance heritability based on our our method in the absence of dominance effect under 16 different genetic architectures. We varied the MAF range of causal variants (MAF of CV), the coupling of MAF with effect size (*a*), and the effect of local LD on effect size (*b* = 0 indicates no LDAK weights and *b* =1 indicates LDAK weights (see Methods). Probability of rejection is computed from 100 replicates. We report p-value of a test of the null hypothesis of no bias in the estimates of 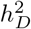 (Methods).

| % of causal SNPs | Genetic architecture | | P(rejection at $p < 0.05$) | Test of bias<br>p-value |
| --- | --- | --- | --- | --- |
|  | MAF of causal SNPs | MAF and LD coupling |  |  |
| 0.01 | [0.01,0.05] | a=b=0 | 6% | 0.192 |
| 0.01 | [0.01,0.05] | a=0,b=1 | 1% | 0.006 |
| 0.01 | [0.01,0.05] | a=0.75,b=0 | 1% | 0.011 |
| 0.01 | [0.01,0.05] | a=0.75,b=1 | 2% | 0.187 |
| 0.01 | [0.0,0.5] | a=b=0 | 1% | 0.388 |
| 0.01 | [0.0,0.5] | a=0,b=1 | 6% | 0.415 |
| 0.01 | [0.0,0.5] | a=0.75,b=0 | 4% | 0.593 |
| 0.01 | [0.0,0.5] | a=0.75,b=1 | 1% | 0.367 |
| 0.01 | [0.05,0.5] | a=b=0 | 3% | 0.046 |
| 0.01 | [0.05,0.5] | a=0,b=1 | 0% | 0.813 |
| 0.01 | [0.05,0.5] | a=0.75,b=0 | 6% | 0.105 |
| 0.01 | [0.05,0.5] | a=0.75,b=1 | 5% | 0.855 |
| 1.0 | [0.0,0.5] | a=b=0 | 1% | 0.196 |
| 1.0 | [0.0,0.5] | a=0,b=1 | 3% | 0.298 |
| 1.0 | [0.0,0.5] | a=0.75,b=0 | 4% | 0.522 |
| 1.0 | [0.0,0.5] | a=0.75,b=1 | 1% | 0.130 |

Next, we considered simulations under genetic architectures with a non-zero dominance effect. We evaluated the accuracy of additive and dominance heritability estimates across 16 different genetic architecture where we vary the additive and dominance heritabilities and proportion of causal dominance variants (Methods). We ran our method using 24 bins for additive effects (based on 6 MAF and 4 LD bins) and a single bin for dominance effects. We obtained accurate estimates of 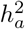 and 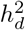 when we jointly fit additive and dominance heritability : biases range from −1.6 × 10^−3^ to 2.7 × 10^−4^ where 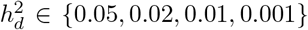 for dominance heritability while the biases range from −2.3 × 10^−3^ to 1.4 × 10^−4^ where 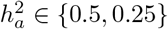 for additive heritability (Figure 2).

**Figure 2:**
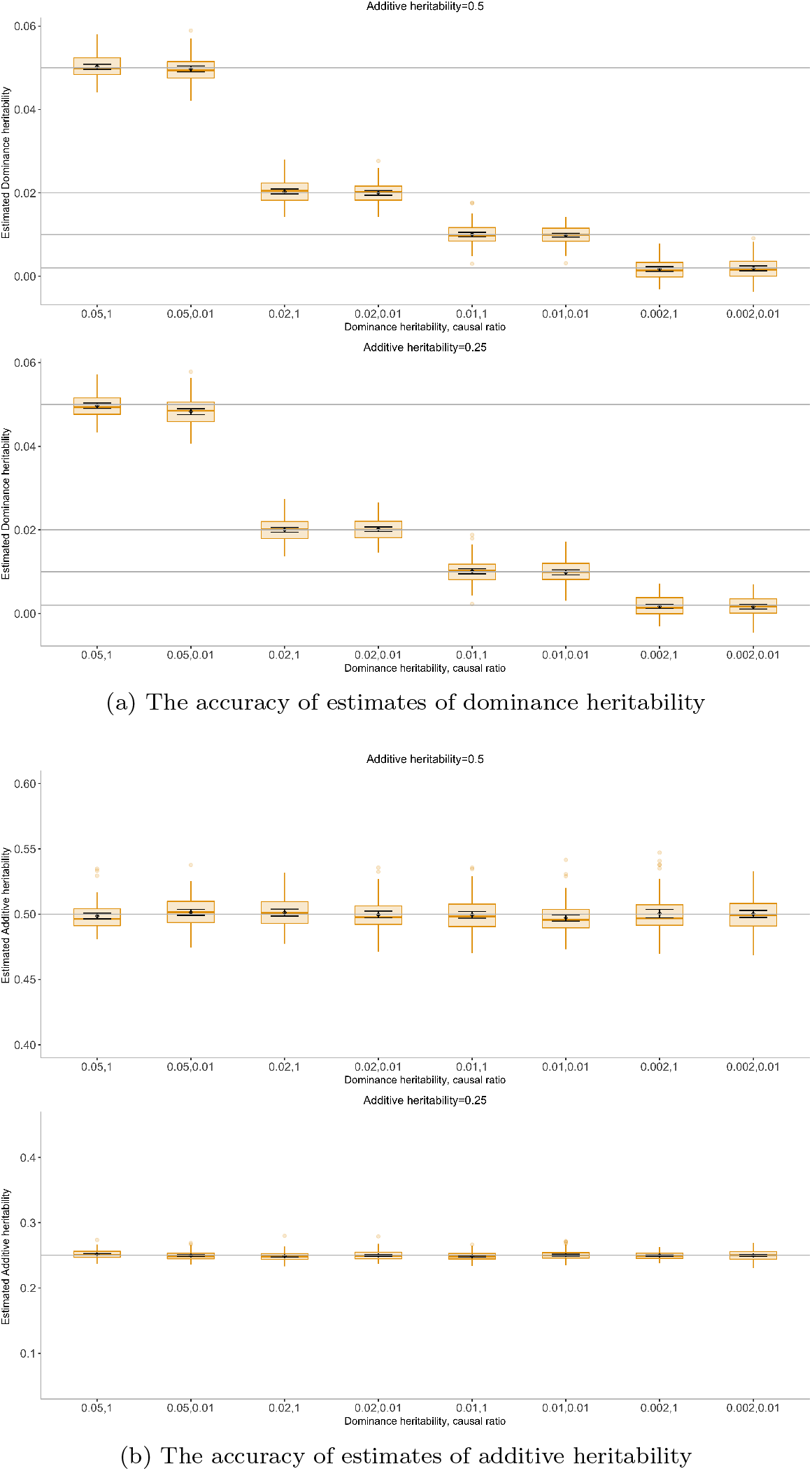
The accuracy of estimates of dominance and additive heritabilities in simulations with non-zero dominance heritability (*N* = 291, 273 unrelated individuals, *M* = 459, 792 array SNPs). In **a**, **b**: We plot estimates from our method under 16 different genetic architectures. We varied the additive heritability 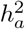, dominance heritability 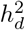, and the proportion of dominance causal variants (causal ratio) (see Methods). Black points and error bars represent the mean and ±2 SE. Each boxplot represents estimates from 100 simulations.

In addition, we observe high power (> 95% for a p-value threshold of 0.05) to detect dominance heritability as low as 1% in a sample size of ≈ 300, 000 (Table 2). A more realistic assessment of power would consider the multiple testing burden incurred when testing a collection of phenotypes with the goal of discovering traits with significant dominance heritability. Assuming we test fifty phenotypes (matching our analyses of the UK Biobank), we estimate 100% power to detect 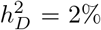 and > 50% power to detect 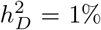 in a sample of ≈ 300,000 individuals 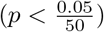.

**Table 2:** Accuracy and power to detect dominance heritability in simulations (*N* = 291, 273 unrelated individuals, *M* = 459, 792 array SNPs). We assess power, bias, and SE of our method in the presence of dominance and additive heritability under 16 different genetic architectures. Power, mean and SE are computed from 100 replicates. We report p-value of a test of the null hypothesis of no bias in the estimates of 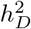 (Methods). Here, *p_causal_*(*A*) and *p_causal_*(*D*) denote proportion of additive and dominance causal variants respectively. 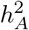 and 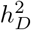 denotes total additive and dominance heritabilities. For both components we assumed GCTA model which is defined as setting *a* = *b* = 0 in Equation 9. Power is reported for p-value threshold of *t* ∈ {0.05, 0.001}.

| Genetic architecture | | Power | | $\hat{h}_D^2$ | | Test of bias |
| --- | --- | --- | --- | --- | --- | --- |
| Additive component | Dominance component | $t = 0.05$ | $t = 10^{-3}$ | Mean | SE | p-value |
| $p_{causal}(A) = 1, h_A^2 = 0.5$ | $p_{causal}(D) = 1, h_D^2 = 0.05$ | 100% | 100% | 0.05 | 0.003 | 0.432 |
| $p_{causal}(A) = 1, h_A^2 = 0.5$ | $p_{causal}(D) = 0.01, h_D^2 = 0.05$ | 100% | 100% | 0.049 | 0.003 | 0.596 |
| $p_{causal}(A) = 1, h_A^2 = 0.5$ | $p_{causal}(D) = 1, h_D^2 = 0.02$ | 100% | 100% | 0.02 | 0.002 | 0.351 |
| $p_{causal}(A) = 1, h_A^2 = 0.5$ | $p_{causal}(D) = 0.01, h_D^2 = 0.02$ | 100% | 100% | 0.02 | 0.002 | 0.869 |
| $p_{causal}(A) = 1, h_A^2 = 0.5$ | $p_{causal}(D) = 1, h_D^2 = 0.01$ | 97% | 76% | 0.01 | 0.002 | 0.901 |
| $p_{causal}(A) = 1, h_A^2 = 0.5$ | $p_{causal}(D) = 0.01, h_D^2 = 0.01$ | 98% | 90% | 0.0099 | 0.002 | 0.730 |
| $p_{causal}(A) = 1, h_A^2 = 0.5$ | $p_{causal}(D) = 1, h_D^2 = 0.002$ | 11% | 0% | 0.0018 | 0.0025 | 0.738 |
| $p_{causal}(A) = 1, h_A^2 = 0.5$ | $p_{causal}(D) = 0.01, h_D^2 = 0.002$ | 10% | 0% | 0.0019 | 0.0027 | 0.590 |
| $p_{causal}(A) = 1, h_A^2 = 0.25$ | $p_{causal}(D) = 1, h_D^2 = 0.05$ | 100% | 100% | 0.049 | 0.003 | 0.434 |
| $p_{causal}(A) = 1, h_A^2 = 0.25$ | $p_{causal}(D) = 0.01, h_D^2 = 0.05$ | 100% | 100% | 0.048 | 0.003 | 2.5e-06 |
| $p_{causal}(A) = 1, h_A^2 = 0.25$ | $p_{causal}(D) = 1, h_D^2 = 0.02$ | 100% | 100% | 0.02 | 0.002 | 0.889 |
| $p_{causal}(A) = 1, h_A^2 = 0.25$ | $p_{causal}(D) = 0.01, h_D^2 = 0.02$ | 100% | 100% | 0.02 | 0.002 | 0.476 |
| $p_{causal}(A) = 1, h_A^2 = 0.25$ | $p_{causal}(D) = 1, h_D^2 = 0.01$ | 93% | 56% | 0.01 | 0.002 | 0.744 |
| $p_{causal}(A) = 1, h_A^2 = 0.25$ | $p_{causal}(D) = 0.01, h_D^2 = 0.01$ | 90% | 61% | 0.0098 | 0.002 | 0.632 |
| $p_{causal}(A) = 1, h_A^2 = 0.25$ | $p_{causal}(D) = 1, h_D^2 = 0.002$ | 8% | 0% | 0.0017 | 0.0024 | 0.373 |
| $p_{causal}(A) = 1, h_A^2 = 0.25$ | $p_{causal}(D) = 0.01, h_D^2 = 0.002$ | 7% | 0% | 0.0017 | 0.0026 | 0.292 |

### Estimates of additive and dominance effects in the UK Biobank

We applied our method to estimate additive and dominance heritability for 50 quantitative traits in the UK Biobank [18]. We restricted our analysis to *N* = 291, 273 unrelated white British individual and *M* = 459, 792 SNPs (MAF> 1%) that were present in the UK Biobank Axiom array. Further, we chose a subset of 50 traits our of a total of 57 traits that have evidence for non-zero additive heritability (Z-score > 3; Methods).

Across the 50 traits, we observe that the average additive heritability 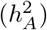 is 21.86% (standard deviation of 9.21% across traits) (Figure 3). On the other hand, we estimate average dominance heritability 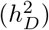 to be 0.13% (standard deviation of 0.39%). On average, we observe that dominance heritability is about 0.48% of additive heritability. We find no evidence for traits that have statistically significant non-zero dominance heritability after correcting for multiple testing.

**Figure 3:**
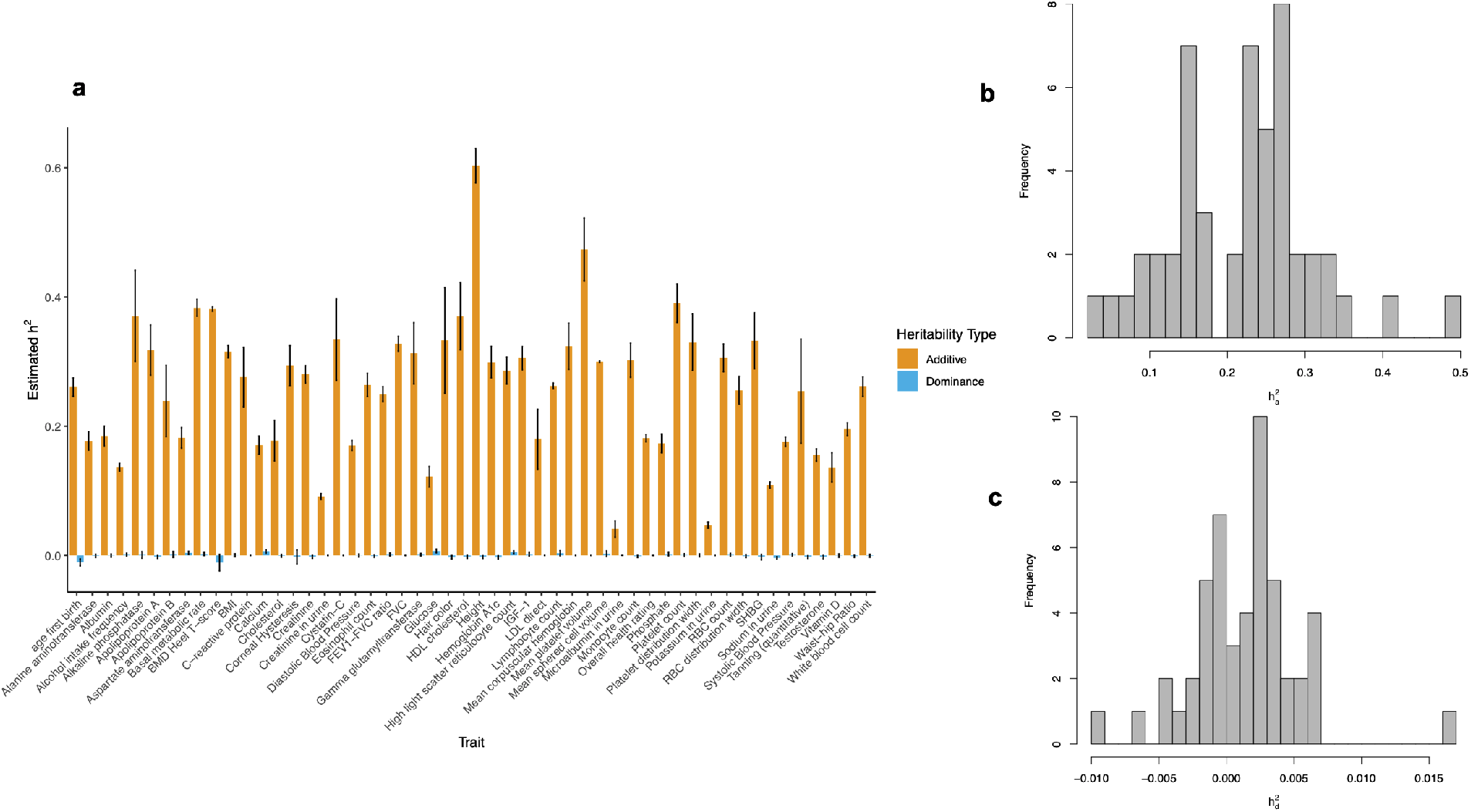
Estimates of additive and dominance heritability for 50 quantitative phenotypes in the UK Biobank (*N* = 291, 273 unrelated white British individuals, *M* = 459, 792 common array SNPs (MAF > 1%)). We ran our method partitioning the additive component into 8 bins defined based on two MAF bins (MAF≤ 0.05, MAF> 0.05) and quartiles of the LD-scores and a single dominance bin. We summarize the estimates of additive and dominance heritability across the 50 phenotypes. In **a** : Black error bars mark ±2 standard errors centered on the estimated heritability. In **b** and **c** we plot the histogram of 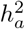 and 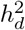 respectively. Point estimates and SE’s are reported in Supplementary Table S2.

It is plausible that our estimates of additive and dominance heritability are underestimated due to imperfect tagging of causal variants by array SNPs. To explore this issue, we analyzed *M* = 4, 824, 392 imputed genotypes with MAF > 1%. Across the 50 traits, we observe that the average 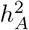 is 22.83% (standard deviation of 9.49% across traits) (Figure 4). The Pearson’s correlation between the point estimates of 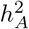 across array and imputed genotypes is 0.998. While the mean ratio of 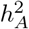 on imputed genotypes to array genotypes being 1.05, we did not observe statistically significant differences between the 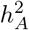 estimates. We estimate 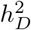 to be 0.06% (standard deviation of 0.19%). On average, we observe that the dominance heritability is about 0.47% of additive heritability. Further, we did not observe any statistically significant differences between the 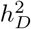 estimates across array and imputed genotypes. We did not find evidence for statistically significant non-zero 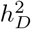 after correcting for multiple testing. We find suggestive evidence for non-zero dominance heritability for blood biochemistry traits: aspartate, basal metabolic rate, blood reticulocyte count, glucose, and calcium (*p* < 0. 05).

**Figure 4:**
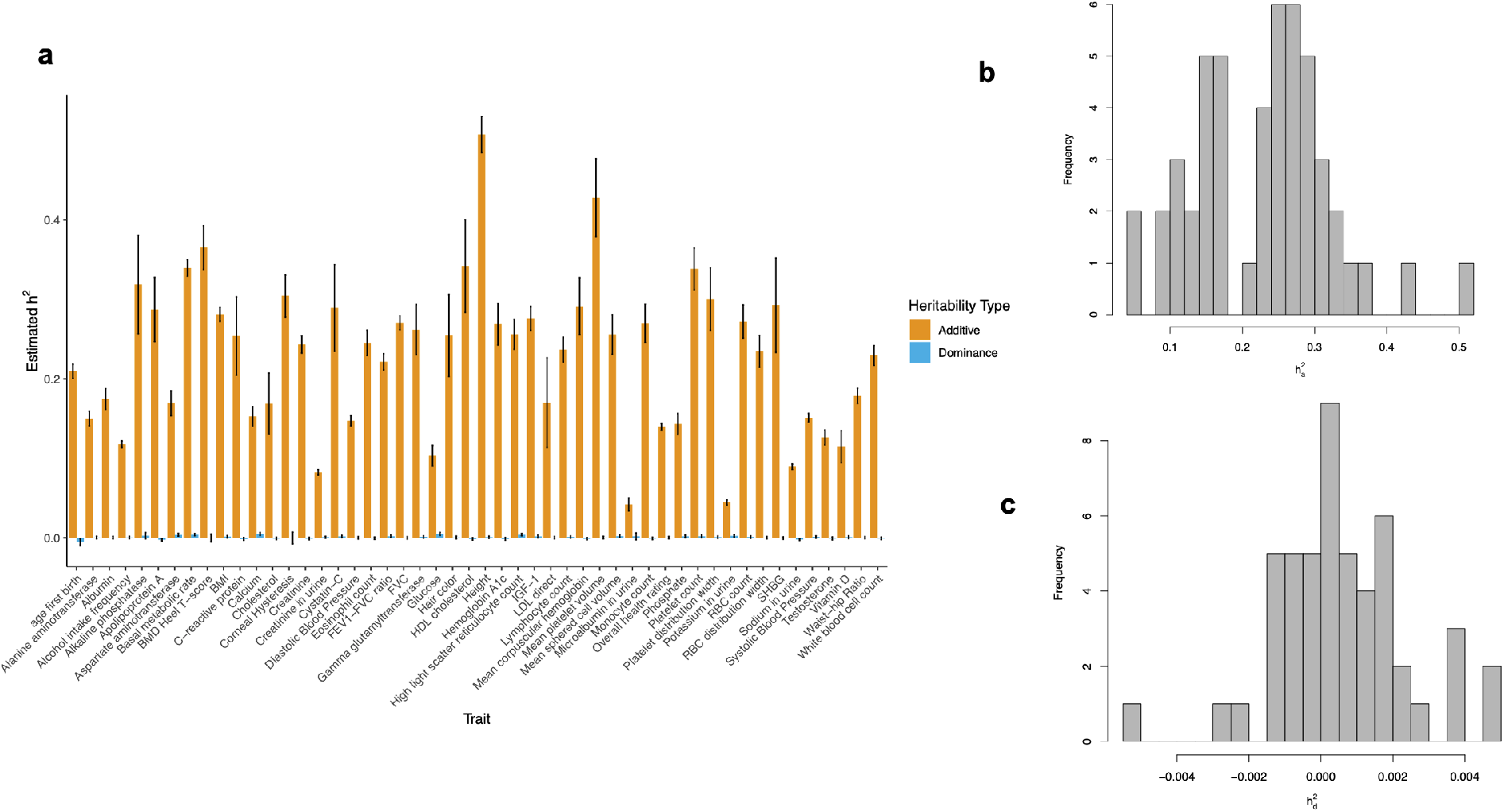
Estimates of additive and dominance heritability for 50 quantitative phenotypes in the UK Biobank (*N* = 291, 273 unrelated white British individuals, *M* = 4, 824, 392 common imputed SNPs (MAF > 1%)). We ran our method partitioning the additive component into 8 bins defined based on two MAF bins (MAF≤ 0.05, MAF> 0.05) and quartiles of the LD-scores and a single dominance bin. We summarize the estimates of additive and dominance heritability across the 50 phenotypes. In **a** : Black error bars mark ±2 standard errors centered on the estimated heritability. In **b** and **c** we plot the histogram of 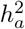 and 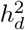 respectively. Point estimates and SE’s are reported in Supplementary Table S3.

## Discussion

The contribution of non-additive genetic effects to complex trait variation has been intensely debated [2, 3, 19, 8, 5]. Here, we have presented a method that can jointly estimate multiple additive and dominance components on biobank-scale genotype-trait data. our method accurately estimates additive and dominance heritability across a range of MAF and LD-dependent genetic architectures. In architectures with no dominance component, tests for the existence of a dominance component based on our method have well-controlled false positive rates. Further, our method has high power to detect dominance components with 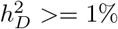 in a sample of ≈ 300*K* unrelated individuals. In application to 50 quantitative traits in the UK Biobank with genotypes measured across 459, 792 common SNPs genotyped on the UK Biobank Axiom array (MAF > 1%) as well as genotypes measured across 4, 824, 392 imputed SNPs (MAF> 1%), we observe substantial additive heritability (21.86% on average for array SNPs, 22.83% on average for imputed SNPs). On the other hand, dominance heritability is substantially lower (0.13% for array and 0.06% for imputed SNPs) and we do not find any trait with statistically significant evidence of dominance heritability. Our power calculations indicate that it is unlikely that 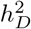 is larger than 1% at the traits analyzed. Taken together, our results suggest that systematic identification of dominance heritability will require analysis of even larger sample sizes than the ≈ 300K individuals that we analyzed here. While the growth of Biobank-scale datasets will facilitate such estimates, such analyses will also require the development of novel methods that can analyze data at scale.

We discuss several limitations of our study as well as directions for future work. The analysis of dominance variance that we have undertaken relies on a specific encoding of dominance and additive effects that leads to uncorrelated components [8]. Alternative encodings might be associated with different statistical and biological interpretation [5]. Second, while our analysis has focused primarily on common SNPs (MAF > 1%), previous work has shown that dominance effects tend to decay faster due to imperfect tagging relative to additive effects leading to a larger bias in estimates of these effects [8]. The concordance of our results across array and imputed genotypes suggests that our estimates of dominance heritability attributed to common SNPs are likely to be robust although we would still underestimate the contribution from low-frequency SNPs. The scalability of our method allows for the exploration of alternative encodings and low-frequency variants at scale. Finally, while our current work focuses on quantitative traits, methods that have previously proposed to estimate heritability in case-control studies [14, 20] can be extended to estimate dominance heritability for binary traits.

## URLs

RHE-mc software: https://github.com/sriramlab/RHE-mc

## Acknowledgments

This research was conducted using the UK Biobank Resource under applications 33127 and 33297. We thank the participants of UK Biobank for making this work possible. This work was funded by NIH grants R01HG009120 (B.P. and K.S.B.), T32HG002536 (A.M.C.), R35GM125055 (S.S.), an Alfred P. Sloan Research Fellowship (S.S.), and NSF grants DGE-1829071 (A.M.C.), III-1705121 (A.P. and S.S.).

## Methods

### Variance components model with additive and dominance components

We aim to fit a variance components model that relates phenotypes ***y*** measured across *N* individuals to their additive and dominance values over *M* SNPs :

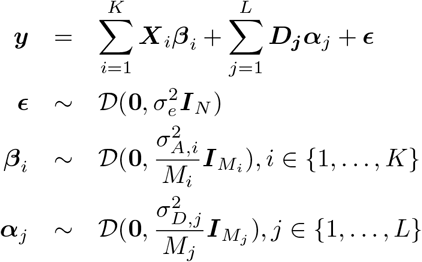

Here 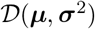 is an arbitrary distribution over a random vector with mean ***μ*** and covariance matrix **Σ**. Here SNPs are partitioned to *K* additive categories *L* dominance categories. Here ***X**_i_* and ***D**_j_* are the *N* × *M_i_* and *N* × *M_j_* matrices consisting of standardized additive and dominance values of SNPs belonging to additive category *i* and dominance category *j* respectively, 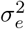 is the residual variance, and 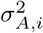 and 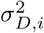 are the variance component of *i*-th additive and dominance categories respectively.

The variance explained by additive variation at all SNPs is defined as:

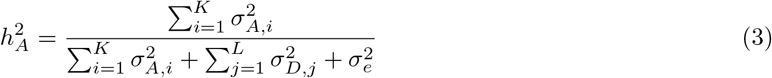

The variance explained by dominance variation at all SNPs is defined as:

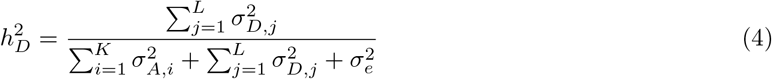

### Method-of-moments for estimating variance components

To estimate the variance components, we use a Method-of-Moments (MoM) estimator that estimates parameter values so that the population moments are close to the sample moments [21]. Since 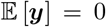, we derived the MoM estimates by equating the population covariance to the empirical covariance. The population covariance is given by:

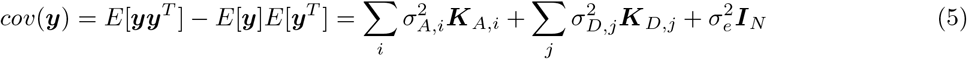

Here 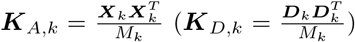 is the additive (dominance) genetic relatedness matrix (GRM) computed from all SNPs of *k*-th category. Using ***yy**^T^* as our estimate of the empirical covariance, we need to solve the following least squares problem to find the variance components.

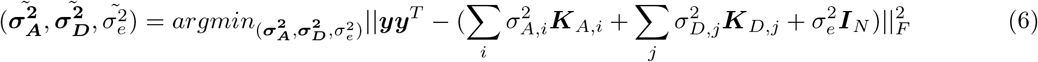

For simplicity, we denote ***K**_i_* = ***K**_A,i_* for *i* = 1,.., *K*,***K**_K+j_* = ***K**_D,j_* for *j* = 1,.., *L* and *J* = *K* + *L*. The MoM estimator satisfies the following normal equations:

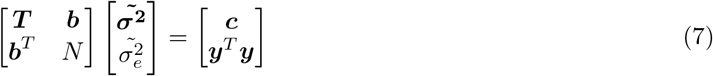

Here 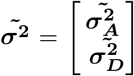, ***T*** is a *J* × *J* matrix with entries *T_k,l_* = *tr*(***K**_k_**K**_l_*), *k, l* ∈ {1,…, *J*}, ***b*** is a *J*-vector with entries *b_k_* = *tr*(***K**_k_*) = *N* (because ***X**_k_^s^* and ***D**_k_^s^* are standardized), and ***c*** is a *J*-vector with entries *c_k_* = ***y**^T^**K**_k_**y***. Each GRM ***K**_k_* can be computed in time 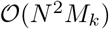 and 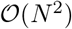 memory. Given *J* GRMs, the quantities *T_k,l_, c_k_, k, l* ∈ {1,…, *J*}, can be computed in 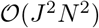. Given the quantities *T_k_,_l_, c_k_*, the normal Equation (7) can be solved in 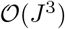. Therefore, the total time complexity for estimating the variance components is 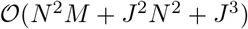

#### Randomized estimator of multiple variance components

The key bottleneck in solving the normal Equation (7) is the computation of *T_k_,_l_, k, l* ∈ {1,…, *J*} which takes 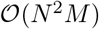. Instead of computing the exact value of *T_k,l_*, we use an unbiased estimator of the trace [22] based on the following identity: for a given *N* × *N* matrix ***C***, ***z**^T^**Cz*** is an unbiased estimator of *tr*(***C***) (*E*[***z**^T^**Cz***] = *tr*[***C***]) where ***z*** be a random vector with mean zero and covariance ***I**_N_*. Hence, we can estimate the values *T_k,l_, k, l* ∈ {1,…, *J*} as follows:

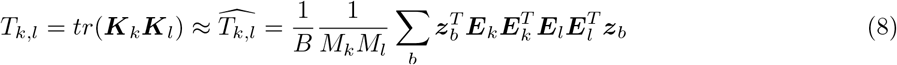

where ***E_i_*** matrix can be standardized additive ***X_i_*** or dominance ***D_i_*** matrix. Here ***z***_i_,…, ***z**_B_* are *B* independent random vectors with zero mean and covariance ***I**_N_*. We draw these random vectors independently from a standard normal distribution. Computing *T_k,l_* using the unbiased estimator involves four multiplications of sub-matrices of the genotype matrix with a vector, repeated *B* times. Therefore, the total running time for estimating the matrix ***T*** is 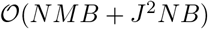.

### Simulations

We simulated phenotypes for UK Biobank genotypes consisting of *M* = 459, 792 array SNPs and *N* = 291, 273 individuals. We obtained the individuals by keeping unrelated white British individuals which are > 3^*rd*^ degree relatives (defined as pairs of individuals with kinship coefficient < 1 /2^(9/2)^)[18], and removing individuals with putative sex chromosome aneuploidy. SNPs were selected as described in the Section on UK Biobank data. We simulated phenotypes from genotypes using the following model:

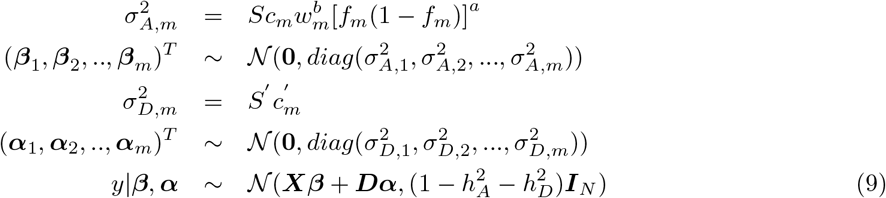

where 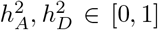, *a* ∈ {0, 0.75}, *b* ∈ {0,1}. Here *S* and *S′* are normalizing constants chosen so that 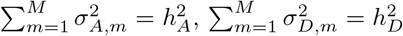. Additive and dominance effect sizes are denoted by ***β*** and ***α*** respectively. *f_m_* and *w_m_* are the minor allele frequency and LDAK score of *m^th^* SNP respectively. In this model, *c_m_*, 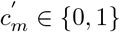 are indicator variables for the causal status of SNP m. The LD score of a SNP is defined to be the sum of the squared correlation of the SNP with all other SNPs that lie within a specific distance, and the LDAK score of a SNP is computed based on local levels of LD such that the LDAK score tends to be higher for SNPs in regions of low LD [23]. The above models relating genotype to phenotype are commonly used in methods for estimating SNP heritability: the GCTA Model (when *a* = *b* = 0 in Equation 9), which is used by the software GCTA [24], and the LDAK Model (where *a* = 0.75, *b* = 1 in Equation (9)) used by software LDAK [23]. Moreover, under each model, we varied the proportion and minor allele frequency (MAF) of causal variants (CVs). Proportion of causal variants were set to be either 100% or 1%, and MAF of causal variants drawn uniformly from [0, 0.5] or [0.01, 0.05] or [0.05, 0.5] to consider genetic architectures that are either infinitesimal or sparse as well genetic architectures that include a mixture of common and rare SNPs as well as one that includes only common SNPs. We generated 100 sets of simulated phenotypes for each setting of parameters.

In experiments to assess the false positive rate, the additive heritability was set to 0.5 while the dominance heritability was set to 0. We computed p-values of a test of the null hypothesis of no 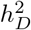 by computing the Z-score of the estimated 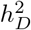 to its standard error and computing a p-value of the two-tailed test. To test the bias of the estimator, for every simulation setting, first we compute 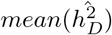 and 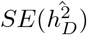 from all replicates, then we reported p-values from Z-score as 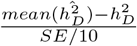.

We also considered a small-scale simulation with *M* = 459, 792 array SNPs and *N* = 10, 000 individuals.

### Power

To assess the power of our method to detect dominance heritability, we considered simulations under different genetic architectures with a non-zero dominance effect. Across 16 different genetic architectures, we vary the additive and dominance heritabilities and proportion of causal dominance variants (Methods). We simulated 100 replicates for every genetic architecture. Let 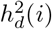 be the estimate of our method on *i*-th replicate for *i* ∈ {1,.., 100}. We computed p-value of *i*-th replicate from Z-score as 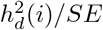 for *i* ∈ {1,.., 100} where SE is computed based on 100 replicates. Finally, we reported the percentage of replicates with p-value< *t* as power of our method on a given simulated genetic architecture for a p-value threshold of *t*.

### UK Biobank Data

In this study, SNPs with greater than 1% missingness and minor allele frequency smaller than 1% were removed. Moreover, SNPs that fail the Hardy-Weinberg test at significance threshold 10^−7^ were removed. We restricted our study to self-reported British white ancestry individuals which are > *3^rd^* degree relatives that is defined as pairs of individuals with kinship coefficient < 1/2^(9/2)^ [18]. Furthermore, we removed individuals who are outliers for genotype heterozygosity and/or missingness. Finally we obtained a set of *N* = 291, 273 individuals and *M* = 459, 792 SNPs to use in the real data analyses. We included age, sex, and the top 20 genetic principal components (PCs) as covariates in our analysis for all traits. We used PCs precomputed by the UK Biobank from a superset of 488, 295 individuals. Additional covariates were used for waist-to-hip ratio (adjusted for BMI) and diastolic/systolic blood pressure (adjusted for cholesterol-lowering medication, blood pressure medication, insulin, hormone replacement therapy, and oral contraceptives).

Further, we analyzed *M* = 4, 824, 392 imputed SNPs with greater than 1% missingness and greater than 1% minor allele frequency. We excluded SNPs that fail the Hardy-Weinberg test at significance threshold 10^−7^. We obtained a set of *N* = 291, 273 individuals and *M* = 4, 824, 392 imputed SNPs.

## Data availability

Access to the UK Biobank resource is available via application at: http://www.ukbiobank.ac.uk.

## Code availability

RHE-mc software is open-source software freely available at: https://github.com/sriramlab/RHE-mc

