## Supplementary Information for "Quantifying the contribution of dominance effects to complex trait variation in biobank-scale data"

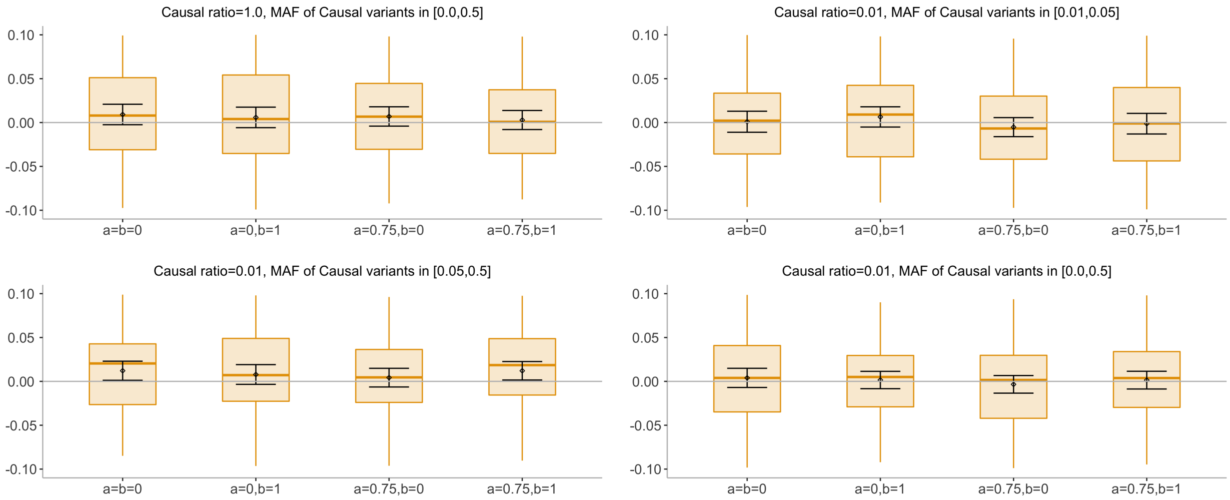


(a) The accuracy of estimates of dominance heritability

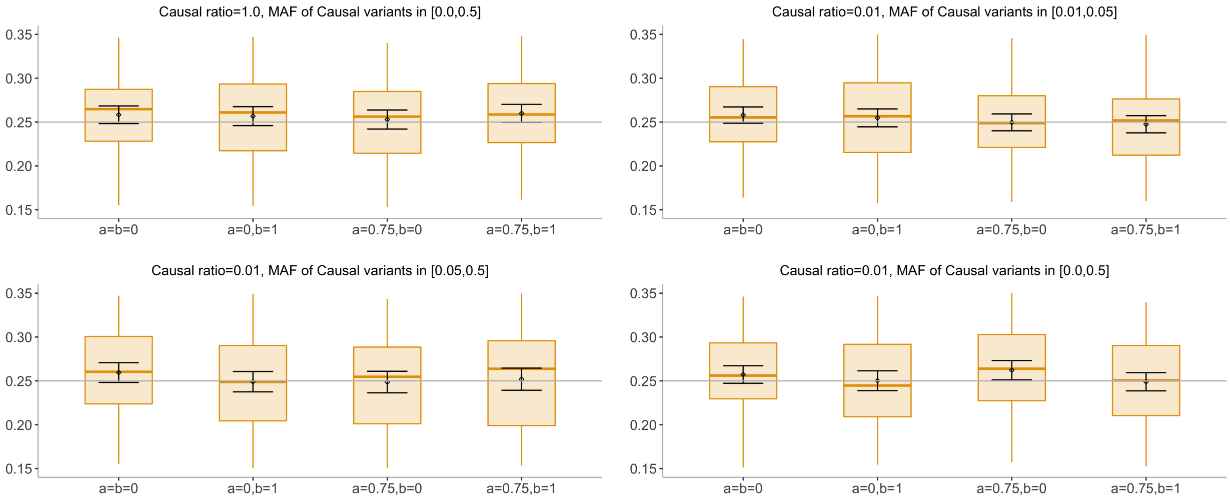


(b) The accuracy of estimates of additive heritability

Figure S1: **The accuracy of estimates of dominance and additive heritabilities in small-scale simulations with no true dominance heritability (**$N = 10,000$ **unrelated individuals,** $M = 459,792$ **array SNPs)**. In **a** and **b**: We plot estimates from our method in the absence of dominance effect under 16 different genetic architectures. We varied the MAF range of causal variants (MAF of CV), the coupling of MAF with effect size ($a$), and the effect of local LD on effect size ($b = 0$ indicates no LDAK weights and $b = 1$ indicates LDAK weights (see Methods). We ran 100 replicates where the true additive and dominance heritabilities of the phenotype are 0.5 and 0.0 respectively. We run our method using single dominance bin and 24 additive bins formed by the combination of 6 bins based on MAF as well as 4 bins based on quartiles of the LDAK score of a SNP (see Methods). Black points and error bars represent the mean and $\pm 2$ SE. Each boxplot represents estimates from 100 simulations.

| Genetic architecture | | | | P(rejection at $p < 0.05$) |
|---|---|---|---|---|
| Percentage of causal SNPs | MAF of causal SNPs | MAF and LD coupling | True $h_D^2$ | |
| 0.01 | [0.01,0.05] | a=b=0 | 0.0 | 2% |
| 0.01 | [0.01,0.05] | a=0,b=1 | 0.0 | 2% |
| 0.01 | [0.01,0.05] | a=0.75,b=0 | 0.0 | 4% |
| 0.01 | [0.01,0.05] | a=0.75,b=1 | 0.0 | 3% |
| 0.01 | [0.0,0.5] | a=b=0 | 0.0 | 2% |
| 0.01 | [0.0,0.5] | a=0,b=1 | 0.0 | 3% |
| 0.01 | [0.0,0.5] | a=0.75,b=0 | 0.0 | 8% |
| 0.01 | [0.0,0.5] | a=0.75,b=1 | 0.0 | 4% |
| 0.01 | [0.05,0.5] | a=b=0 | 0.0 | 7% |
| 0.01 | [0.05,0.5] | a=0,b=1 | 0.0 | 2% |
| 0.01 | [0.05,0.5] | a=0.75,b=0 | 0.0 | 7% |
| 0.01 | [0.05,0.5] | a=0.75,b=1 | 0.0 | 6% |
| 1.0 | [0.0,0.5] | a=b=0 | 0.0 | 2% |
| 1.0 | [0.0,0.5] | a=0,b=1 | 0.0 | 3% |
| 1.0 | [0.0,0.5] | a=0.75,b=0 | 0.0 | 3% |
| 1.0 | [0.0,0.5] | a=0.75,b=1 | 0.0 | 4% |

Table S1: **Calibration of tests of non-zero dominance heritability in small-scale simulations**. We assess calibration of our method in the absence of dominance effect under 16 different genetic architectures. We varied the MAF range of causal variants (MAF of CV), the coupling of MAF with effect size ($a$), and the effect of local LD on effect size ($b = 0$ indicates no LDAK weights and $b = 1$ indicates LDAK weights (see Methods). Probability of rejection is computed from 100 replicates.

| Trait | Heritability | |
|---|---|---|
| | Additive | Dominance |
| Age first birth | $0.20125 \pm 0.00815$ | $-0.00187 \pm 0.00632$ |
| Alanine aminotransferase | $0.14268 \pm 0.00903$ | $-0.00018 \pm 0.00265$ |
| Albumin | $0.16201 \pm 0.01128$ | $0.00333 \pm 0.00283$ |
| Alcohol intake frequency | $0.11322 \pm 0.00415$ | $-0.00197 \pm 0.00235$ |
| Alkaline phosphatase | $0.31743 \pm 0.07676$ | $0.01605 \pm 0.01909$ |
| Apolipoprotein A | $0.27997 \pm 0.03758$ | $-0.00256 \pm 0.00244$ |
| Aspartate aminotransferase | $0.16841 \pm 0.02172$ | $0.00686 \pm 0.00267$ |
| Basal metabolic rate | $0.32489 \pm 0.00868$ | $0.00253 \pm 0.00263$ |
| BMD Heel T-score | $0.34912 \pm 0.02914$ | $-0.00087 \pm 0.00694$ |
| BMI | $0.26371 \pm 0.00874$ | $0.00395 \pm 0.00257$ |
| C-reactive protein | $0.25196 \pm 0.0488$ | $-0.00119 \pm 0.00252$ |
| Calcium | $0.14078 \pm 0.01216$ | $0.00449 \pm 0.0029$ |
| Cholesterol | $0.15793 \pm 0.03074$ | $0.00188 \pm 0.00295$ |
| Corneal Hysteresis | $0.29504 \pm 0.02625$ | $-0.00917 \pm 0.01182$ |
| Creatinine | $0.23893 \pm 0.00985$ | $-0.00178 \pm 0.00225$ |
| Creatinine in urine | $0.07615 \pm 0.00352$ | $0.00175 \pm 0.00234$ |
| Cystatin-C | $0.27628 \pm 0.04872$ | $0.00542 \pm 0.00278$ |
| Diastolic Blood Pressure | $0.14413 \pm 0.00511$ | $0.00181 \pm 0.00239$ |
| Eosinophil count | $0.22676 \pm 0.01476$ | $-4e-04 \pm 0.00262$ |
| FEV1-FVC ratio | $0.21179 \pm 0.01061$ | $0.0027 \pm 0.00263$ |
| FVC | $0.27173 \pm 0.00726$ | $-0.00473 \pm 0.00241$ |
| Gamma glutamyltransferase | $0.2542 \pm 0.03487$ | $0.00222 \pm 0.00259$ |
| Glucose | $0.09337 \pm 0.01063$ | $0.00649 \pm 0.00295$ |
| Hair color | $0.25831 \pm 0.05408$ | $-0.00202 \pm 0.00345$ |
| HDL cholesterol | $0.32793 \pm 0.05241$ | $-9e-04 \pm 0.003$ |
| Height | $0.49507 \pm 0.0192$ | $-0.00045 \pm 0.00285$ |
| Hemoglobin A1c | $0.25886 \pm 0.02513$ | $0.00219 \pm 0.00283$ |
| High light scatter reticulocyte count | $0.23626 \pm 0.01749$ | $0.00215 \pm 0.00265$ |
| IGF-1 | $0.26427 \pm 0.01478$ | $0.00368 \pm 0.00288$ |
| LDL direct | $0.15062 \pm 0.04315$ | $0.00225 \pm 0.00306$ |
| Lymphocyte count | $0.2214 \pm 0.01295$ | $0.00314 \pm 0.00257$ |
| Mean corpuscular hemoglobin | $0.27188 \pm 0.03247$ | $0.00075 \pm 0.00273$ |
| Mean platelet volume | $0.41688 \pm 0.04684$ | $0.00698 \pm 0.00292$ |
| Mean sphered cell volume | $0.23819 \pm 0.02096$ | $0.00607 \pm 0.00254$ |
| Microalbumin in urine | $0.03864 \pm 0.012$ | $-0.00642 \pm 0.00915$ |
| Monocyte count | $0.2569 \pm 0.02192$ | $0.00145 \pm 0.00265$ |
| Overall health rating | $0.14046 \pm 0.00363$ | $-0.00322 \pm 0.00233$ |
| Phosphate | $0.13263 \pm 0.01091$ | $-0.00026 \pm 0.00299$ |
| Platelet count | $0.31935 \pm 0.02465$ | $0.00515 \pm 0.0028$ |
| Platelet distribution width | $0.2824 \pm 0.03317$ | $0.00358 \pm 0.00275$ |
| Potassium in urine | $0.04415 \pm 0.00261$ | $0.0021 \pm 0.00212$ |
| RBC count | $0.26032 \pm 0.0218$ | $0.00232 \pm 0.00296$ |
| RBC distribution width | $0.2216 \pm 0.01688$ | $0.00292 \pm 0.0031$ |
| SHBG | $0.27094 \pm 0.04282$ | $0.00016 \pm 0.0026$ |
| Sodium in urine | $0.08761 \pm 0.00355$ | $-0.00068 \pm 0.00247$ |
| Systolic Blood Pressure | $0.14555 \pm 0.00466$ | $0.00232 \pm 0.00257$ |
| Testosterone | $0.13135 \pm 0.00794$ | $-0.00447 \pm 0.00288$ |
| Vitamin D | $0.10642 \pm 0.01868$ | $0.0049 \pm 0.0024$ |
| Waist-hip Ratio | $0.16782 \pm 0.00813$ | $-0.00173 \pm 0.00232$ |
| White blood cell count | $0.22091 \pm 0.01219$ | $0.00038 \pm 0.00268$ |

Table S2: **Estimate the proportions of variance explained by additive and dominance variation from our method for 50 quantitative phenotypes in the UK Biobank ($N = 291,273$ unrelated white British individuals, $M = 459,792$ common SNPs)**. We run our method partitioning the additive component into 8 bins defined based on two MAF bins (MAF$\leq 0.05$, MAF$> 0.05$) and quartiles of the LD-scores and a single dominance bin.

| Trait | Heritability | |
|---|---|---|
| | Additive | Dominance |
| Age first birth | $0.20954 \pm 0.00893$ | $-0.00519 \pm 0.00402$ |
| Alanine aminotransferase | $0.15026 \pm 0.00933$ | $0.00037 \pm 0.00163$ |
| Albumin | $0.17477 \pm 0.01302$ | $0.00026 \pm 0.00157$ |
| Alcohol intake frequency | $0.11771 \pm 0.00436$ | $0.00042 \pm 0.00155$ |
| Alkaline phosphatase | $0.31876 \pm 0.06188$ | $0.00265 \pm 0.00399$ |
| Apolipoprotein A | $0.28728 \pm 0.04033$ | $-0.00254 \pm 0.00147$ |
| Aspartate aminotransferase | $0.1693 \pm 0.01576$ | $0.00379 \pm 0.00177$ |
| Basal metabolic rate | $0.3396 \pm 0.0104$ | $0.00398 \pm 0.00138$ |
| BMD Heel T-score | $0.36517 \pm 0.02759$ | $-0.00018 \pm 0.0049$ |
| BMI | $0.28143 \pm 0.0089$ | $0.00142 \pm 0.00151$ |
| C-reactive protein | $0.25389 \pm 0.04917$ | $-0.00143 \pm 0.00176$ |
| Calcium | $0.15274 \pm 0.01237$ | $0.0048 \pm 0.00183$ |
| Cholesterol | $0.16889 \pm 0.03858$ | $-0.00087 \pm 0.00153$ |
| Corneal Hysteresis | $0.30444 \pm 0.02675$ | $-0.00028 \pm 0.00743$ |
| Creatinine | $0.24341 \pm 0.01095$ | $-0.001 \pm 0.00156$ |
| Creatinine in urine | $0.08243 \pm 0.00365$ | $0.00078 \pm 0.0012$ |
| Cystatin-C | $0.28918 \pm 0.05443$ | $0.00168 \pm 0.00152$ |
| Diastolic Blood Pressure | $0.14725 \pm 0.00659$ | $-5e-05 \pm 0.00153$ |
| Eosinophil count | $0.24544 \pm 0.01602$ | $-0.00063 \pm 0.00149$ |
| FEV1-FVC ratio | $0.22161 \pm 0.01022$ | $0.00183 \pm 0.00161$ |
| FVC | $0.27073 \pm 0.00878$ | $-0.00011 \pm 0.00154$ |
| Gamma glutamyltransferase | $0.26184 \pm 0.03161$ | $0.00108 \pm 0.00147$ |
| Glucose | $0.10327 \pm 0.01324$ | $0.0049 \pm 0.002$ |
| Hair color | $0.25466 \pm 0.05171$ | $0.00042 \pm 0.00181$ |
| HDL cholesterol | $0.3419 \pm 0.05838$ | $-0.00141 \pm 0.0015$ |
| Height | $0.50749 \pm 0.0229$ | $0.00079 \pm 0.00141$ |
| Hemoglobin A1c | $0.26891 \pm 0.02624$ | $-0.00138 \pm 0.00176$ |
| High light scatter reticulocyte count | $0.25615 \pm 0.01887$ | $0.00397 \pm 0.00147$ |
| IGF-1 | $0.27632 \pm 0.01504$ | $0.00169 \pm 0.00171$ |
| LDL direct | $0.17004 \pm 0.05658$ | $0.00015 \pm 0.00152$ |
| Lymphocyte count | $0.23694 \pm 0.0155$ | $0.00101 \pm 0.00136$ |
| Mean corpuscular hemoglobin | $0.29132 \pm 0.036$ | $-0.00114 \pm 0.00142$ |
| Mean platelet volume | $0.42801 \pm 0.0491$ | $0.00034 \pm 0.00162$ |
| Mean sphered cell volume | $0.25593 \pm 0.02494$ | $0.00206 \pm 0.00159$ |
| Microalbumin in urine | $0.04201 \pm 0.0081$ | $0.00181 \pm 0.00441$ |
| Monocyte count | $0.26996 \pm 0.02427$ | $-0.00086 \pm 0.00152$ |
| Overall health rating | $0.13942 \pm 0.00426$ | $0.00021 \pm 0.00147$ |
| Phosphate | $0.14328 \pm 0.01327$ | $0.00176 \pm 0.00169$ |
| Platelet count | $0.33832 \pm 0.0261$ | $0.00184 \pm 0.00171$ |
| Platelet distribution width | $0.30071 \pm 0.03937$ | $8e-04 \pm 0.00149$ |
| Potassium in urine | $0.04449 \pm 0.0032$ | $0.00239 \pm 0.00147$ |
| RBC count | $0.27218 \pm 0.0208$ | $0.00097 \pm 0.00153$ |
| RBC distribution width | $0.23468 \pm 0.01997$ | $0.00012 \pm 0.00146$ |
| SHBG | $0.29273 \pm 0.05944$ | $-0.00036 \pm 0.00156$ |
| Sodium in urine | $0.08935 \pm 0.00354$ | $-0.00217 \pm 0.00125$ |
| Systolic Blood Pressure | $0.15109 \pm 0.00554$ | $0.00107 \pm 0.00142$ |
| Testosterone | $0.12642 \pm 0.00934$ | $-0.00082 \pm 0.00186$ |
| Vitamin D | $0.11467 \pm 0.02034$ | $0.00079 \pm 0.0017$ |
| Waist-hip Ratio | $0.17866 \pm 0.00949$ | $0.00019 \pm 0.00135$ |
| White blood cell count | $0.22953 \pm 0.01254$ | $-0.00078 \pm 0.00158$ |

Table S3: **Estimate the proportions of variance explained by additive and dominance variation from our method for 50 quantitative phenotypes in the UK Biobank ($N = 291,273$ unrelated white British individuals, $M = 4,824,392$ common SNPs).** We run our method partitioning the additive component into 8 bins defined based on two MAF bins (MAF$\leq 0.01$, MAF$> 0.01$) and quartiles of the LD-scores and a single dominance bin.
